# Hydroxychloroquine and Zinc ameliorate interleukin-6 associated hepato-renal toxicity induced by *Aspergillus fumigatus* in experimental models

**DOI:** 10.1101/2023.09.05.556428

**Authors:** Jude Ogechukwu Okoye, Anslem Tochukwu Basil, Onyedikachi Okoli, Precious Onyemaechi Achebe

## Abstract

In Nigeria, immunocompromised persons, particularly those living with HIV, are at an increased risk of developing invasive pulmonary aspergillosis caused by *Aspergillus fumigatus*. Interestingly, this condition produces symptoms that can be easily mistaken for those of COVID-19. To better understand the pathophysiology of Aspergillosis and determine the therapeutic and toxic effects of Zinc and HCQ, this study examined liver and renal functions in experimental models. This experimental study included 28 Albino rats, assigned into 7 Groups (n= 4 each); designated A to G. Group A received the standardized rat chow and distilled water only. Group B received a moderate dose of HCQ only. Group C received *A. fumigatus* suspension (AFS) without any treatments. Group D simultaneously received AFS and a low dose of HCQ. Group E simultaneously received AFS and a moderate dose of HCQ. Group F simultaneously received AFS and a high dose of HCQ. Group G simultaneously received AFS and a moderate dose of HCQ and Zinc. Serum levels of interleukins (IL)-6 and 10, liver enzymes, and renal parameters were measured accordingly. The lungs, liver, and kidneys were excited and weighed. Significance was set at p< 0.05. Higher levels of serum alanine transaminase, creatinine, and urea and lower relative lung weight were observed in group C compared with other groups (p< 0.001). Higher IL-6 levels and IL-6/IL-10 ratio were also observed in group C compared with other groups (p> 0.05). In conclusion, this study revealed that HCQ and Zinc ameliorate oxidative stress and damage induced by *A. fumigatus*.

## Introduction

Aspergilli are filamentous molds that may lead to a wide spectrum of infections including, hypersensitivity and allergic diseases depending on the host’s immune status or pulmonary structure, chronic pulmonary infections, and acute life-threatening infections [1]. Aspergillosis is caused by a type of mold called Aspergillus fumigatus. Pulmonary aspergillosis is a collective term used to refer to several conditions caused by infection with a fungus of Aspergillus species, especially Aspergillus fumigatus affecting the respiratory tracts basically the Lungs [2]. On rare occasions, this pathogen can disseminate to other organs, such as the brain and kidneys. An invasive variant of Aspergillosis is a devastating illness, with mortality rates in some patient groups reaching as high as 90% [3]. Over the previous two decades, the incidence of invasive aspergillosis has increased tenfold [4]. Aspergillosis is basically seen in immunocompromised patients and can also be symptomatic in certain cases. A prevalence of 3.1 % was detected among individuals living with HIV in Nigeria [5]. Studies have shown that Hydroxychloroquine (HCQ) modulates the immune system by interfering with lysosomal acidification inhibition of antigen presentation, and down-regulation of cytokine production and secretion by monocytes and T cells [6]. Hydroxychloroquine inhibits cytokine production and modulation of certain co-stimulatory molecules [7] while Zinc is an essential vital exogenous mineral that aids DNA synthesis, enzymatic reactions, immune functions, protein synthesis, wound healing, growth, and development [8]. Inflammatory cytokines such as interleukin (IL)-6 and IL-10 are important biomarkers for distinguishing infections caused by various pathogens [9]. IL-6 and Interferon gamma are noted to be predominantly elevated in invasive pulmonary aspergillosis and Pneumocystis pneumonia [9]. IL-10 increases the host’s susceptibility to lethal fungal infection, possibly because IL-10 is associated with a Th2 response, down-regulation of a Th1 response, and macrophage activation [10]. With the rising frequency of pulmonary aspergillosis and diagnosis in Nigeria, chronic obstructive pulmonary disease, it has become critical to assess the efficacy of HCQ and zinc in alleviating respiratory distress associated with pulmonary aspergillosis. For the first time, this study investigated the curative and toxic effects of different doses of HCQ and zinc on experimentally induced respiratory distress following exposure to *Aspergillus fumigatus*.

## Methods

### Study site

The ethical approval for this research was obtained from the College of Health Sciences Ethics Committee at Nnamdi Azikiwe University. The animals were carefully handled following ethical standards and protocols for the care and use of laboratory animals by the National Institute of Health in 2011. This experimental study was carried out at the vivarium in the Department of Physiology. Nnamdi Azikiwe University, Nnewi Campus, Nigeria.

### Collection of isolate and induction of Aspergillosis

The *Aspergillus fumigatus* isolate was obtained from a patient’s sample in Onitsha and sub-cultured on Saboraud dextrose agar (SDA) at 37°C for 2 days. The conidia were extracted by washing the plates with sterile 0.2% Tween 20 and Norma saline, followed by centrifugation and filtration of the suspension. The animals were subjugated to anesthetics to make them unconscious via the use of the anesthetic “Ketamine”. This drug (0.5ml of ketamine) was then administered slowly (over a period of 60 seconds). The conidia suspension was administered into the right nasal cavity via the use of a tuberculin syringe. The rat nostrils pointed facing upwards and suspension of *Aspergillus fumigatus* conidia dropped at a slow rate (150ul) into the right nasal cavity with the tuberculin string gently inserted 1.5cm deep into the nasal cavity. On both days 8, 9, and 10 animals were also exposed to A. fumigatus again via inhalation of the conidia spores.

### Determination of ED50 (Median Effective Dose)

The oral LD50 of HCQ in rats is 1240 mg/kg, while the therapeutic dose is 10mg/kg (MSDS, 2008 cited in [11] and the oral lethal dose 50 (LD50) for Zinc salts in rats is 237–623 mg/kg [12], and LD50 of Hydrocortisone acetate is 150mg/kg. The intraperitoneal LD50 of cyclophosphamide without halothane was 251 mg/kg, and with 2 hours subsequent exposure to 152 mg/kg of halothane [13].

### Study design and animal handling

The test system for this study was Albino rats (N= 28), weighing 80LJ±LJ20 g (Table 1). The animals were allowed to acclimatize in the vivarium for two weeks prior to the start of the experiment. They were randomly divided into seven groups, each containing four animals. Groups C to G were given a moderate dose of steroid-sparing agent (Cyclophosphamide 75mg/kg, ∼0.3ml intraperitoneally) on day 1 and a steroid (Hydrocortisone acetate 80mg/kg, ∼0.32ml intraperitoneally) with the aim of reducing the inflammatory response, replicating a real-life situation as Aspergillosis majorly affects immunosuppressed individuals, thus favouring the development of the disease. The steroids were administered orally using an oral cannula and the oral cannula also applies to the HCQ and Zinc. On day 6, Aspergillosis was sufficiently introduced using Aspergillosis suspension in drops into the right nasal cavity using tuberculin string, with the fact already established that, 5 days after the intraperitoneal injection of Cyclophosphamide (steroid-sparing agent) 75mg/kg and (a steroid) Hydrocortisone acetate (80mg/kg) that the WBC’s count would be sufficiently decreased [14]. On day 7, a second batch of immunosuppressants was administered (CPA intraperitoneally at a dose of 60mg/kg) to ensure immunosuppression. Starting from day 13 to day 15, the respective animals were given their corresponding doses of Hydroxychloroquine and zinc.

**Table 1:**
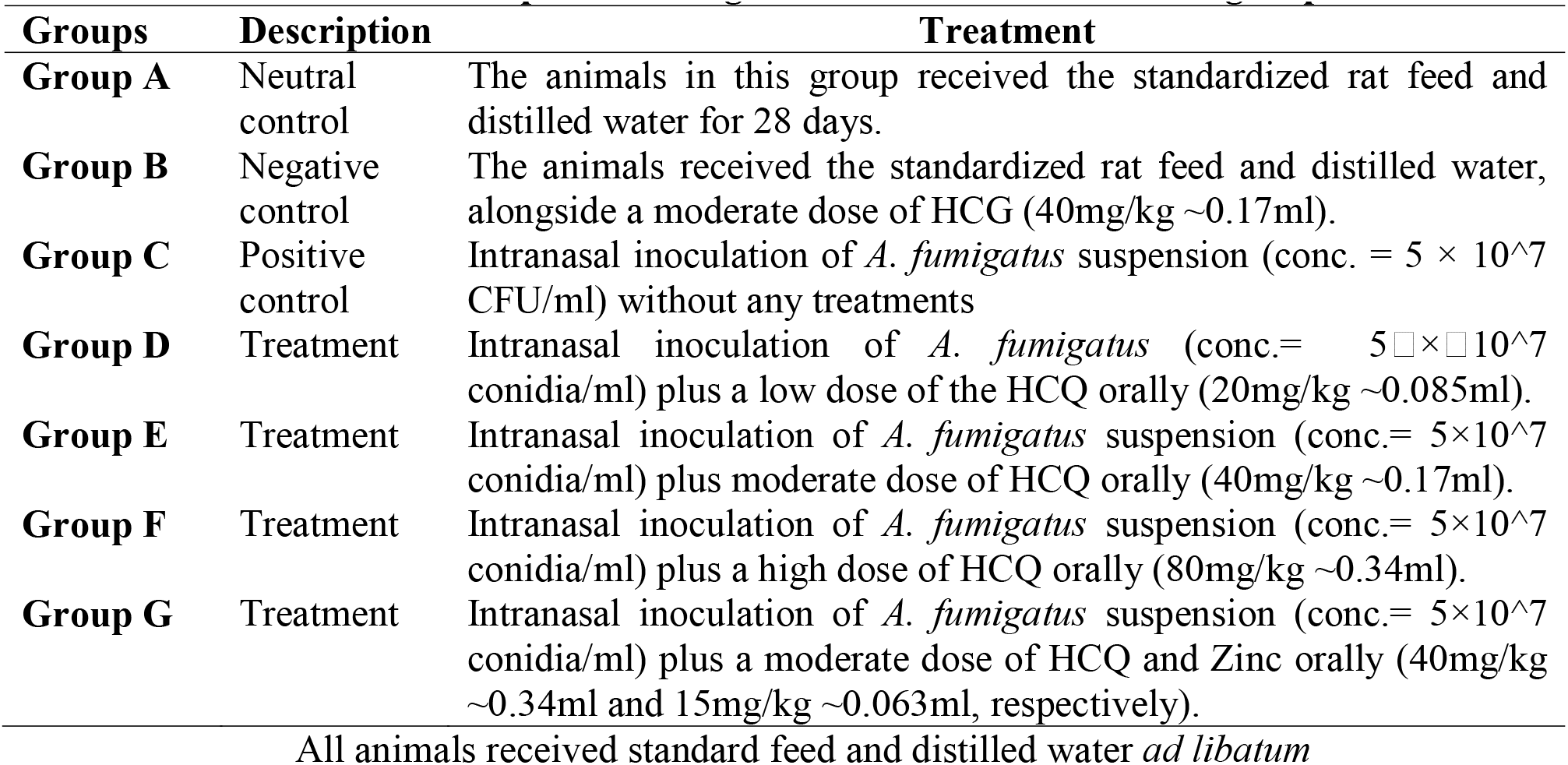
Description and regimen for the control and test groups.

| Groups | Description | Treatment |
| --- | --- | --- |
| **Group A** | Neutral control | The animals in this group received the standardized rat feed and distilled water for 28 days. |
| **Group B** | Negative control | The animals received the standardized rat feed and distilled water, alongside a moderate dose of HCG (40mg/kg ~0.17ml). |
| **Group C** | Positive control | Intranasal inoculation of *A. fumigatus* suspension (conc. = $5 \times 10^7$ CFU/ml) without any treatments |
| **Group D** | Treatment | Intranasal inoculation of *A. fumigatus* (conc.= $5 \times 10^7$ conidia/ml) plus a low dose of the HCQ orally (20mg/kg ~0.085ml). |
| **Group E** | Treatment | Intranasal inoculation of *A. fumigatus* suspension (conc.= $5 \times 10^7$ conidia/ml) plus moderate dose of HCQ orally (40mg/kg ~0.17ml). |
| **Group F** | Treatment | Intranasal inoculation of *A. fumigatus* suspension (conc.= $5 \times 10^7$ conidia/ml) plus a high dose of HCQ orally (80mg/kg ~0.34ml). |
| **Group G** | Treatment | Intranasal inoculation of *A. fumigatus* suspension (conc.= $5 \times 10^7$ conidia/ml) plus a moderate dose of HCQ and Zinc orally (40mg/kg ~0.34ml and 15mg/kg ~0.063ml, respectively). |

All animals received standard feed and distilled water *ad libitum*

On day 16, the animals were made to fast, and the sacrifice was carried out on day 17 (the 31^st^ day after acclimatization).

### Sample Collection and Laboratory Assessment

On the 31^st^ day, the animals were anesthetized by placing the animals in an air-tight jar containing cotton wool soaked in chloroform. Following loss of reflexes, response to stimulus, and reduction in the animal’s respiratory rate, blood samples were collected by ocular puncture. The blood samples were collected in EDTA anticoagulant bottles (Alpha Surgicare) and plain bottles, labeled, and left undisturbed for 15 minutes. The whole blood samples were centrifuged for 10 minutes at 3000 rpm to separate the serum. The lungs, kidneys, and liver were subsequently harvested, processed, and stained using the Haematoxylin and Eosin staining technique.

### Determination of Interleukins 6 and 10 levels

To evaluate cytokine storm and interleukin (IL) drift, levels of IL-6 and IL-10 were carried out using the ELISA Microwell Test Kit. The density of yellow is proportional to the target amount of sample captured in the plate. Read the O.D. absorbance at 450nm in a microplate reader, and then the concentration of the target can be calculated. Before proceeding with the assay, all reagents, serum, calibrators, and controls were brought to room temperature (20-27°C) and the test Procedure was carefully carried out. The IL-6 ELISA kit Fine (Test ER0042) possessed a reactivity range of 62.5 - 4000pg/ml and a sensitivity of 37.5pg/ml, while the IL-10 ELISA test kit Fine (Test ER0033) possessed a reactivity of 31.25 - 2000pg/ml and a sensitivity of 18.75pg/ml. The reference range for IL-6 was 0 to 43.5pg/ml.

### Statistical Analysis

Statistical Science for Social Sciences (SPSS) Version 25 was used to analyze the data obtained from this work. Relative organ weight was calculated as the weight of the organ (Lungs, Liver, and Kidney) divided by total body weight multiplied by 100. One-way Analysis of Variance (ANOVA) was used to analyze data from the biochemical assay as well as relative organ weights (organ weight over total body weight multiplied by 100). The *Post Hoc* test was used for group comparison. To determine the relationships between parameters, Pearson’s correlation was used. Data were considered significant at *p* ≤ 0.05, ≤ 0.01, and ≤ 0.001.

## Results

### General Observation

All the experimental rats in groups C, D, E, F, and G had decreased appetite after immune inhibitor injection. The animals subsequently started exhibiting lethargy and a decrease in their reflexes four (4) days after immune suppression and recovery nine days after immune suppression. On the second day after the first fungal inoculation, sneezing and nose scratching were observed in the Aspergillosis-induced groups and increased with subsequent fungal inoculations.

### Interleukin 6 and 10 (IL-6 and IL-10) levels and ratio

The pro-inflammatory IL-6 was higher in group C compared with other groups at p> 0.05 (Table 2). Hydroxychloroquine caused a substantial decrease in IL-6 (in group B) and with increasing dosage of HCQ (towards the LD-50), an increase in IL-6 was also noted due to oxidative stress. A higher IL-10 was observed in group G compared with other groups. This suggests that a combination of HCQ and Zinc mitigates the inflammation induced by Aspergillosis. In recent years, the ratio of proinflammatory cytokine (IL-6) to anti-inflammatory cytokine (IL-10), (IL-6/IL-10 ratio) has been used as a reliable marker for measuring inflammatory status. There were dose-dependent differences in IL-6 and IL-10 levels among animals treated with low, moderate, and high doses of HCQ. The results also show that group C had the worst outcome compared with other treatment groups (p> 0.05) due to the higher IL6/IL-10 ratio in the group (Figure 1). There was a direct relationship between IL-6 and IL-10 (r= 0.649, p< 0.001).

**Table 2:** Mean comparison of interleukins and liver enzyme levels across experimental groups.

| Groups | IL-6 | IL-10 | AST | ALT | ALP |
| --- | --- | --- | --- | --- | --- |
| | Mean ± SD | Mean ± SD | Mean ± SD | Mean ± SD | Mean ± SD |
| **Group A** | 4.41 ± 0.78 | 112.92 ± 24.04 | 10.50 ± 1.29 | 9.50 ± 2.38 | 54.25 ± 19.52 |
| **Group B** | 4.58 ± 0.30 | 109.34 ± 14.82 | 13.75 ± 1.26 | 12.50 ± 0.58 | 71.75 ± 19.52 |
| **Group C** | 5.11 ± 0.36 | 115.37 ± 12.30 | 12.50 ± 4.51 | 15.75 ± 2.50 | 104.50 ± 33.73 |
| **Group D** | 3.61 ± 1.18 | 101.23 ± 24.39 | 9.75 ± 2.06 | 8.75 ± 0.96 | 81.50 ± 21.30 |
| **Group E** | 3.87 ± 0.56 | 102.24 ± 10.94 | 10.25 ± 0.96 | 10.00 ± 2.31 | 65.50 ± 9.88 |
| **Group F** | 3.89 ± 0.51 | 116.3 ± 3.63 | 9.75 ± 2.06 | 9.75 ± 1.71 | 80.00 ± 23.51 |
| **Group G** | 4.88 ± 1.40 | 127.92 ± 27.26 | 11.00 ± 1.15 | 9.50 ± 1.29 | 64.75 ± 15.73 |
| F-value | 1.870 | 0.969 | 1.864 | 7.422 | 2.258 |
| P-value | 0.134 | 0.470 | 0.135 | 0.000* | 0.077 |
| **A vs B** | 0.777 | 0.788 | 0.050a | 0.029a | 0.264 |
| **A vs C** | 0.248 | 0.854 | 0.215 | 0.000c | 0.003b |
| **A vs G** | 0.432 | 0.267 | 0.753 | 1.000 | 0.498 |
| **B vs C** | 0.378 | 0.652 | 0.434 | 0.019a | 0.043a |
| **B vs D** | 0.110 | 0.544 | 0.018a | 0.008b | 0.529 |
| **B vs E** | 0.234 | 0.595 | 0.036a | 0.065 | 0.686 |
| **B vs F** | 0.247 | 0.602 | 0.018a | 0.044a | 0.594 |
| **B vs G** | 0.612 | 0.172 | 0.094 | 0.029a | 0.651 |
| **C vs D** | 0.018a | 0.294 | 0.094 | 0.094 | 0.146 |
| **C vs E** | 0.046a | 0.330 | 0.166 | 0.166 | 0.018a |
| **C vs F** | 0.049a | 0.944 | 0.094 | 0.094 | 0.123 |
| **C vs G** | 0.704 | 0.350 | 0.349 | 0.349 | 0.016a |
| **D vs G** | 0.041a | 0.055 | 0.434 | 0.434 | 0.284 |

Statistical analysis: ANOVA and Post hoc Test. \*Significance is set at $p \leq 0.05 = a$ , $\leq 0.01 = b$ , $\leq 0.001 = c$ Number of animals per group = 4.

**Figure 1:**
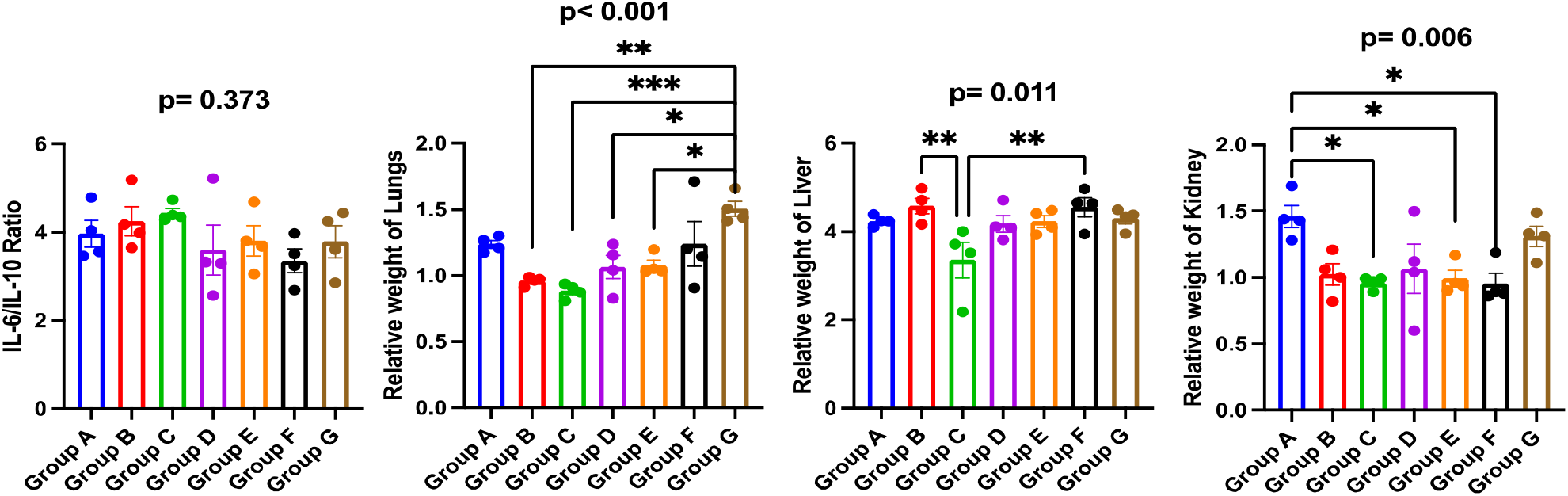
Relative organ weight and Interleukin ratio across experimental groups.

### Dysregulation of liver enzymes

There was a direct relationship between AST and ALT (r= 0.565, p= 0.002), AST and ALP (r= 0.391, p= 0.044), and ALT and ALP (r= 0.588, p= 0.001). A higher AST was observed in group B, treated with HCQ alone, compared with the neutral control (group A) at p< 0.05 (Table 2). This suggests that HCQ is capable of inducing hepatoxicity in animals that are without any underlying disease. However, treatment with HCQ among animals with Aspergillosis resulted in a reduced AST level compared with the infected and untreated groups (p> 0.05). In group B, a high level of ALT was also observed compared with other groups (p< 0.05). Higher ALT and ALP levels were observed in group C, infected *A. fumigatus*, and untreated, compared with other controls and treatment groups at p< 0.05 and p> 0.05, respectively. Infected animals treated with moderate doses of HCQ and Zinc had lower levels of ALP compared with other groups, except for group A (p> 0.05). This also suggests that the addition of Zinc to the treatment regime mitigated the inflammation induced by Aspergillosis and moderate doses of HCQ.

In Figure I, group C had a higher IL-6/IL-10 ratio and lower relative weight of the lungs, liver, and kidneys (consistent with atrophy) compared with other controls and treatment groups. The IL-6/IL-10 and relative organ weight of Group G were like those of Group A. This suggests that the addition of Zinc to a moderate dose of HCQ improved treatment outcome.

### Alterations in renal function

Higher levels of potassium, bicarbonate, creatinine, and urea and lower levels of sodium and chloride were observed in group C compared with other treatment groups at p> 0.05, > 0.05, < 0.05, < 0.05, > 0.05 and > 0.05, respectively (Table 3). A dose-dependent increase in potassium levels was also observed among animals treated with low, moderate, and high doses of HCQ (p> 0.05). Animals that received a moderate dose of HCQ had better renal function compared to animals that received low or high doses of HCQ. Results indicate that adding zinc to a moderate dose of HCQ did not improve renal function compared to treatment with HCQ alone. Pearson’s correlation revealed a direct relationship was observed between potassium and Urea (r= 0.481, p= 0.011), sodium and chloride (r= 0.864, p< 0.001), and Urea and Creatinine (r= 0.700, p< 0.001).

**Table 3:** Mean comparison of renal function parameters across the experiment groups.

| Groups | Potassium | Sodium | Chloride | Bicarbonate | Creatinine | Urea |
| --- | --- | --- | --- | --- | --- | --- |
| | Mean ± SD | Mean ± SD | Mean ± SD | Mean ± SD | Mean ± SD | Mean ± SD |
| **Group A** | 4.83 ± 0.25 | 141.75 ± 3.30 | 100.50 ± 4.12 | 24.25 ± 1.26 | 9.25 ± 1.26 | 42.00 ± 8.76 |
| **Group B** | 4.40 ± 0.59 | 142.00 ± 1.41 | 103.25 ± 0.96 | 23.25 ± 1.71 | 9.75 ± 0.50 | 44.75 ± 4.79 |
| **Group C** | 5.18 ± 0.83 | 141.50 ± 2.38 | 101.25 ± 1.50 | 25.00 ± 0.82 | 14.75 ± 2.22 | 132.25 ± 34.33 |
| **Group D** | 4.28 ± 0.34 | 145.00 ± 1.41 | 104.00 ± 1.41 | 25.00 ± 0.96 | 8.50 ± 1.29 | 49.00 ± 2.83 |
| **Group E** | 4.63 ± 0.43 | 142.50 ± 2.65 | 102.75 ± 2.50 | 23.75 ± 1.71 | 9.50 ± 0.58 | 44.25 ± 6.50 |
| **Group F** | 4.80 ± 0.64 | 143.25 ± 3.10 | 103.75 ± 2.06 | 24.50 ± 1.29 | 8.75 ± 0.96 | 45.50 ± 9.25 |
| **Group G** | 4.95 ± 0.39 | 142.50 ± 1.73 | 102.50 ± 1.29 | 24.75 ± 0.96 | 8.75 ± 1.50 | 49.25 ± 10.01 |
| F-value | 1.388 | 0.985 | 1.354 | 1.201 | 11.252 | 19.873 |
| P-value | 0.265 | 0.460 | 0.278 | 0.344 | 0.000* | 0.000* |
| **A vs C** | 0.362 | 0.884 | 0.637 | 0.419 | 0.000c | 0.000c |
| **A vs D** | 0.158 | 0.069 | 0.036a | 0.284 | 0.425 | 0.508 |
| **A vs F** | 0.948 | 0.386 | 0.050a | 0.786 | 0.594 | 0.740 |
| **B vs C** | 0.052a | 0.771 | 0.215 | 0.068 | 0.000c | 0.000c |
| **B vs D** | 0.742 | 0.091 | 0.637 | 0.039a | 0.190 | 0.687 |
| **C vs D** | 0.026a | 0.051a | 0.094 | 0.786 | 0.000c | 0.000c |
| **C vs E** | 0.158 | 0.561 | 0.349 | 0.184 | 0.000c | 0.000c |
| **C vs F** | 0.329 | 0.313 | 0.125 | 0.588 | 0.000c | 0.000c |
| **C vs G** | 0.555 | 0.561 | 0.434 | 0.786 | 0.000c | 0.000c |

Statistical analysis: ANOVA and Post hoc Test. \*Significance is set at $p \leq 0.05 = a, \leq 0.01 = b, \leq 0.001 = c$ Number of animals per group = 4.

## DISCUSSION

This study assessed the immunological and biochemical parameters effects of HCQ and zinc in experimentally induced invasive pulmonary Aspergillosis. In the study, models with Aspergillosis without any treatments had the highest IL-6 level and low glomerular filtration rate due to renal atrophy. A subsequent decrease in IL-6 was noted in group D, E, and F, where both Aspergillosis invasion was conjugated with varying dose of HCQ, and this was possibly due to the immunosuppressive effects of HCQ. Due to this immunosuppressive and immunomodulatory effect of HCQ, it could be used in the management of many inflammatory diseases. Inflammatory cytokines, especially IL-6, are important biomarkers for distinguishing infection-associated trauma or inflammation caused by various pathogens such as Aspergillosis [9,15,16]. Some cytokines such as IL-10 exert anti-inflammatory and immunosuppressive effects to restore homeostasis [17]. The pro-inflammatory process (primarily IL-6) is subsequently followed by anti-inflammatory (with IL-10 as one of the potent anti-inflammatory cytokines) [15], but the compensatory increase in IL-10 levels is ineffective in counteracting the increasing levels of IL-6 [18]. The IL-6/IL-10 ratio was higher in groups B and C causing them to have a low organ weight and body mass in relationship to the other groups. An increase in the IL-6/IL-10 ratio is directly associated with a poor outcome [19].

The IL-6/IL-10 ratio decreased significantly in group F, which correlated with an increase in oxidative stress and organ damage. After careful analysis, it becomes apparent that there was a reduction in the IL-6 value, but an increase in the IL-10 value in Groups D and group E. This elevation in IL-10 serves to suppress immune responses, warding off autoimmunity, and decreasing oxidative stress levels [20]. Upon the administration of Zinc as an immune booster (Group G), there was an increase in the level of IL-6, irrespective of the synergized effects of the HCQ (which acted as an immunosuppressant-in-Aspergillosis-induced Groups). Zinc induces monocytes to produce interleukin-1, interleukin-6, and tumor necrosis factor-α [21]. Similarly, decreased production of TH1 cytokines and interferon-α by leukocytes in healthy elderly persons is correlated with low zinc serum levels (Ibid). It is important to note that in response to zinc supplementation, plasma cytokines exhibit a dose-dependent response [22].

According to the findings of this study, the use of HCQ in the absence of Aspergillosis resulted in an augmentation of organ weight and elevated liver enzymes. The shift in liver function could potentially be attributed to heightened levels of IL-6. Research indicates that IL-6 may substantially impact B cell hyperactivity and could even directly impact tissue harm [23]. However, results show that HCQ was able to ameliorate Aspergillosis-induced hepatoxicity through concomitant reduction of IL-6 levels and decreased polymorphonuclear infiltrates [24]. Although zinc supplement has not been shown to result in higher expression of IL-10 [25], the results from this study revealed higher expression of IL-10 when HCQ and zinc are simultaneously administered to experimental models. Although HCQ has shown promise in treating respiratory diseases, there are concerns about its potential side effects such as renal toxicity and heart failure, particularly when taken at high doses, as noted by Alanagreh et al. [26]. Nevertheless, the finding of this study suggests that a low dose of HCQ or a combination of a moderate dose of HCQ and zinc may improve renal function when dealing with Aspergillosis. This finding is reinforced by a prior study that highlights the potential beneficial effects of HCQ on renal dysfunction, as outlined by Bourke [27].

## Conclusion

The objective of this study was to explore the effects of HCQ and zinc on Aspergillosis, along with the cytokine storm that comes with it. The research revealed that both HCQ and zinc have the potential to alleviate cytokine levels and relative organ weight. This indicates that HCQ might be able to lower the risk of autoimmunity and systemic inflammatory syndromes that can be life-threatening.

## References

1. Jenks, Jeffrey D. and Hoenigl, M. (2018). Treatment of Aspergillosis. Journal of Fungi, 4(3), pp. 260–263.

2. Gaillard, F. (2021). Pulmonary Aspergillosis | Radiology Reference Article | Radiopaedia.org. [online] Radiopaedia. Available at: https://radiopaedia.org/articles/pulmonary-Aspergillosis/ [Accessed 28 Sep. 2021].

3. Dagenais, T.R.T. and Keller, N.P. (2009). Pathogenesis of Aspergillus fumigatus in Invasive Aspergillosis. Clinical Microbiology Reviews, [online] 22(3), 447–465.

4. Zakaria, A. and Hamze, M. (2020). Recent trends in the epidemiology, diagnosis, treatment, and mechanisms of resistance in clinical Aspergillus species: A general review with a special focus on the Middle Eastern and North African region. Journal of Infection and Public Health, 13(1), 1–10.

5. Adekola, Hafeez A., Agbede, Olajide O., Abdullahi, Idris N., Olayemi, Lawal O., Emeribe, Anthony U., Dangana, A. and Yunusa, T. (2019). Assessment of serum Aspergillus galactomannan expression and associated risk factors among patients living with HIV at Ilorin, Nigeria. Microbiologia Medica, [online] 34(2), 139–167.

6. Varisli, L., Cen, O., and Vlahopoulos, S. (2020). Dissecting pharmacological effects of chloroquine in cancer treatment: interference with inflammatory signaling pathways. Immunology, 159(3), 257–278.

7. Schrezenmeier, E. and Dörner, T. (2020). Mechanisms of action of hydroxychloroquine and chloroquine: implications for rheumatology. Nature Reviews. Rheumatology, [online] 16(3), 155–166.

8. Kubala, J. (2018). Zinc: Benefits, Deficiency, Food Sources and Side Effects. [online] Healthline. Available at: https://www.healthline.com/nutrition/zinc [Accessed 28 Oct. 2021].

9. Shen, H.-P., Tang, Y.-M., Song, H., Xu, W.-Q., Yang, S.-L. and Xu, X.-J. (2016). Efficiency of interleukin 6 and interferon gamma in the differentiation of invasive pulmonary Aspergillosis and pneumocystis pneumonia in pediatric oncology patients. International Journal of Infectious Diseases, 48(3), 73–77.

10. Clemons, K.V., Grunig, G., Sobel, R.A., Mirels, L.F., Rennick, D.M. and Stevens, D.A. (2000). Role of IL-10 in invasive Aspergillosis: increased resistance of IL-10 gene knockout mice to lethal systemic Aspergillosis. Clinical & Experimental Immunology, 122(2), pp. 186–191.

11. ElShishtawy, M., Elgendy, F. and Ramzy, R. (2013). Comparative Toxicity Study of Chloroquine and Hydroxychloroquine on Adult Albino Rats. Mansoura Journal of Forensic Medicine and Clinical Toxicology, 21(2), 15–28.

12. Andriollo-Sanchez, M., Claeyssen, R., Arnaud, J., Touvard, L., Denis, J., Chancerelle, Y., Roussel, A.-M. and Agay, D. (2008). Toxic Effects of Iterative Intraperitoneal Administration of Zinc Gluconate in Rats. Basic & Clinical Pharmacology & Toxicology, 103(3), 267–272.

13. Taniguchi, T., Koido, Y., Aiboshi, J., Yamashita, T., Suzaki, S. and Kurokawa, A. (1999). Change in the ratio of interleukin-6 to interleukin-10 predicts a poor outcome in patients with systemic inflammatory response syndrome. Critical Care Medicine, 27(7), 1262–1264.

14. Yan, Y., Zhao, Z., Wan, H., Wu, R., Fang, J. and Liu, H. (2014). A novel fungus concentration-dependent rat model for acute invasive fungal rhinosinusitis: an experimental study. BioMed Central Infectious Diseases, 14(1), 89–112.

15. Sapan, H.B., Paturusi, I., Islam, A.A., Yusuf, I., Patellongi, I., Massi, M.N., Pusponegoro, A.D., Arief, S.K., Labeda, I., Rendy, L. and Hatta, M. (2017). Interleukin-6 and interleukin-10 plasma levels and mRNA expression in polytrauma patients. Chinese Journal of Traumatology, 20(6), 318–322.

16. Chandler, L.C., Yusuf, I.H., McClements, M.E., Barnard, A.R., MacLaren, R.E. and Xue, K. (2020). Immunomodulatory Effects of Hydroxychloroquine and Chloroquine in Viral Infections and Their Potential Application in Retinal Gene Therapy. International Journal of Molecular Sciences, 21(14), 149–151.

17. Liang, N., Zhong, Y., Zhou, J., Liu, B., Lu, R., Guan, Y., Wang, Q., Liang, C., He, Y., Zhou, Y., Song, J. and Zhou, J. (2018). Immunosuppressive effects of hydroxychloroquine and artemisinin combination therapy via the nuclear factor-κB signaling pathway in lupus nephritis mice. Experimental and Therapeutic Medicine, [online] 15(3), 2436–2442.

18. Rong, Y.-D., Bian, A.-L., Hu, H.-Y., Ma, Y. and Zhou, X.-Z. (2018). Study on relationship between elderly sarcopenia and inflammatory cytokine IL-6, anti-inflammatory cytokine IL-10. BioMed Central Geriatrics, 18(1), 193–220.

19. Angelakis, E., Million, M., Kankoe, S., Lagier, J.-C., Armougom, F., Giorgi, R. and Raoult, D. (2014). Abnormal Weight Gain and Gut Microbiota Modifications Are Side Effects of Long-Term Doxycycline and Hydroxychloroquine Treatment. Antimicrobial Agents and Chemotherapy, 58(6), 3342–3347.

20. Chatzopoulos, G., Doufexi, A.-E., Wolff, L. and Kouvatsi, A. (2018). Interleukin-6 and interleukin-10 gene polymorphisms and the risk of further periodontal disease progression. Brazilian Oral Research, 32(0), 253–289.

21. Rink, L. and Kirchner, H. (2000). Zinc-Altered Immune Function and Cytokine Production. The Journal of Nutrition, 130(5), 1407.

22. Foster, M. and Samman, S. (2012). Zinc and Regulation of Inflammatory Cytokines: Implications for Cardiometabolic Disease. Nutrients, 4(7), pp. 676LJ694.

23. Tackey, E., Lipsky, P. E., & Illei, G. G. (2004). Rationale for interleukin-6 blockade in systemic lupus erythematosus. Lupus, 13(5), 339–343.

24. Alhazzani, K., Alrewily, S.Q., Aljerian, K., Alhosaini, K., Algahtani, M.M., Almutery, M.F., Alhamed, A.S., Nadeem, A., Alotaibi, M.R. and Alanazi, A.Z., 2023. Hydroxychloroquine ameliorates dasatinib-induced liver injury via decrease in hepatic lymphocytes infiltration. Human & Experimental Toxicology, 42, p. 09603271231188492.

25. Rosenkranz, E., Metz, C.H., Maywald, M., Hilgers, R.D., Weßels, I., Senff, T., Haase, H., Jäger, M., Ott, M., Aspinall, R. and Plümäkers, B. (2016). Zinc supplementation induces regulatory T cells by inhibition of SirtLJ1 deacetylase in mixed lymphocyte cultures. Molecular nutrition & food research, 60(3), pp. 661–671.

26. Alanagreh, L. A., Alzoughool, F., & Atoum, M. (2020). Risk of using hydroxychloroquine as a treatment of COVID-19. International Journal of Risk & Safety in Medicine, 31(3), 111–116.

27. Bourke, L., McCormick, J., Taylor, V., Pericleous, C., Blanchet, B., Costedoat-Chalumeau, N., … & Ioannou, Y. (2015). Hydroxychloroquine protects against cardiac ischaemia/reperfusion injury in vivo via enhancement of ERK1/2 phosphorylation. PLoS One, 10(12), e0143771.

